# Electro-composting: an emerging technology

**DOI:** 10.1101/2025.01.16.633320

**Authors:** Ahmad Shabir Hozad, Christian Abendroth

## Abstract

The present study focuses on electrical stimulation for composting. Using the PSALSAR method, a systematic review resulted in 24 relevant articles. For each system, the review describes key materials, composter design, operating conditions, temperature evolution, compost maturity, microbial community, and outcome. The examined studies fall into four main systems: electric field-assisted aerobic composting (EAAC), electrolytic oxygen aerobic composting (EOAC), microbial fuel cells (MFC), and thermoelectric generators (TEG). Apart from the main systems highlighted above, there is a rare case, which remains hardly studied. This includes bioelectrochemically assisted anaerobic composting (AnC_BE, III_). EAAC and EOAC systems biologically balance microbial activity and organic matter decomposition, whereas MFC and TEG systems have dual functioning due to energy generated alongside waste degradation. These innovative systems significantly improve composting efficiency by speeding up organic matter breakdown, increasing oxygen supply, and facilitating energy recovery. Together, they overcome the drawbacks of conventional composting systems and promote more environmentally friendly waste management solutions.

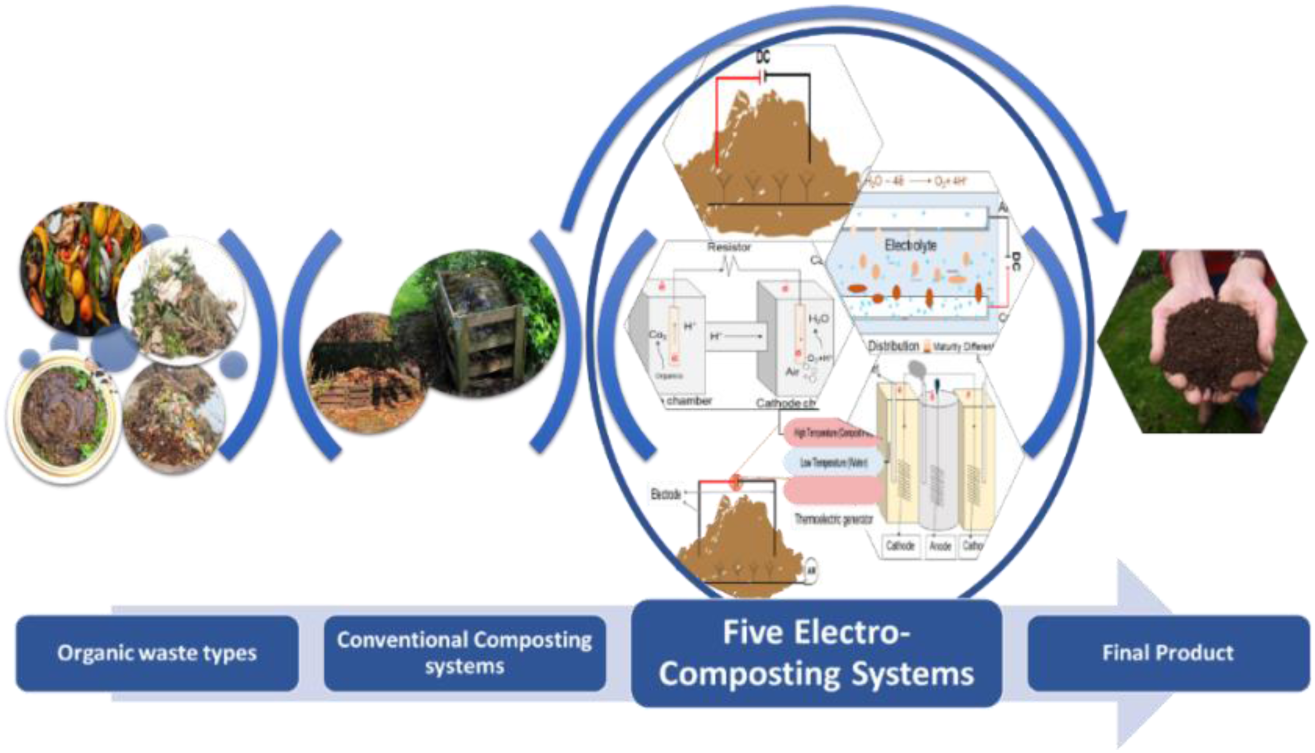

**Research Highlights:**

1. The four most frequently investigated electro-composting systems are EAAC, EOAC, MFC, and TEG.
2. MFCs make it easier to turn organic waste into electricity, which makes decomposition more environmentally friendly.
3. Thermal energy generated during decomposition is utilized by TEGs to produce electricity.
4. The research on AnC_BE_ is highly limited.

## 1. Introduction

Electro-composting has garnered significant attention as an advanced technique in the sustainable management of organic waste, aiming to enhance the efficiency of composting while simultaneously addressing environmental concerns such as greenhouse gas (GHG) emissions and nutrient recycling. Conventional aerobic composting, which is frequently employed for treating organic waste, has several disadvantages, including high GHG emissions, extended maturation times, and low oxygen utilization efficiency (Tang et al., 2019). These challenges have facilitated the development of technologies such as electro-composting, which enhances microbial activity and improves composting conditions by employing electric fields or electrochemical processes (Cao et al., 2021b).

Electric-field assisted aerobic composting (EAAC) is a promising electro-composting system that employs a direct current to elevate compost temperatures, improve microbial activity, and reduce the length of the composting process (Cao et al., 2021a, 2021b, 2022; Tang et al., 2019, 2020). Studies have demonstrated that EAAC systems can increase compost temperatures by 5-10 °C in comparison to conventional methods, thereby improving both oxygen utilization and compost maturity (Fu et al., 2022b) (Fu et al., 2022b). Additionally, electrolytic oxygen aerobic composting (EOAC) employs electrolysis to produce oxygen in situ, a process that has been demonstrated to enhance the degradation of organic material by reducing the formation of anaerobic zones and maintaining aerobic conditions within the compost pile (Fu et al., 2022).

Microbial fuel cells (MFCs) have also developed as a dual-purpose technology that generates bioelectricity and manages wastes. The system provides a novel approach to waste management and energy recovery by utilizing microbial metabolism to convert organic materials into electrical energy (Pant et al., 2010). MFCs have been examined for their capacity to process a diverse array of organic wastes, including rice hulls, oil cake (from mustard plants), leaf mold, grass trimmings, and chicken feces, and generate electricity as a byproduct (Moqsud et al., 2013).

Thermoelectric generators (TEGs) are an additional distinctive method of electro-decomposition that utilizes the heat produced during the composting process to generate electricity (Shangguan et al., 2020).

This review systematically evaluates the most prominent electro-composting systems (EAAC, EOAC, MFC, and TEG) in terms of their environmental impact, operating efficiencies, and technological advancements. Electrical composting is an emerging field and little has been published on this so far. Especially bioelectrochemically assisted anaerobic composting (AnC_BE,III_) is a rare case that has received little attention so far. Despite their lack of attention, these systems offer innovative insights and potential advancements in the field of electro-composting, particularly in the optimization of waste treatment and energy recovery processes. The authors aspire to offer a comprehensive evaluation of the role of electro-composting in the promotion of sustainable waste management methods by contrasting existing systems and highlighting future innovations.

## 2. Material and Methods

The PSALSAR technique (Mengist et al., 2020) was rigorously employed in this systematic literature study to investigate the integration of electrical technology in composting. Initially, a protocol was developed to determine the research topic, objectives, and scope, focusing on the impact of electrical stimulation technologies on composting processes. Boolean operators were employed to identify key search phrases, particularly “compost* AND electri*” or “compost* AND electro*” which were then applied to the Web of Science database. A vast amount of literature (7791 items in total) appeared and many articles were unrelated to the topic. Therefore, a solution was searched for to make the search more specific. This has been done by skimming through the first 250 articles. The recurrent terms and keywords associated with electrical stimulation and composting were examined in these papers. A preliminary list was created by compiling important terms from different parts of these articles including titles, abstracts, keywords, and main sections. 17 particular search terms, such as compost* AND electrode, compost* AND “electric field”, compost* AND MFC, and compost* AND “thermoelectric composting” were chosen following a comprehensive cross-comparison and repetition removal process. The number of articles was reduced to 984 articles after applying these 17 keywords. The outcomes for each search term are represented in Table 1. All Articles were sorted to retain only the article type of “original research” and non-English publications were excluded as well. The remaining 403 articles were screened in more detail, which involved two stages: an initial title and abstract screening that omitted 312 irrelevant articles, and a comprehensive examination of the remaining articles. In this regard, an extensive analysis was conducted to classify all articles into composting processes (91 articles) and articles that employed electrical technologies in combination with composting (22 articles). Table 1 illustrates the number of articles obtained per search term before applying the criterion, while Figures 1 and 2 illustrate the screening process, publication trends, and abundant keywords. Additionally, Figure 3C shows a word cloud that graphically depicts the key terms used in the 24 chosen articles. The abstract and keywords of these articles served as input data for the WordClouds.com software (accessed at https://www.wordclouds.com). The most common terms were “electric composting,” and “aerobic composting,” which appeared more frequently. The size of each term in the word cloud indicates the number of times it is used in the text being analyzed.

**Table 1:**
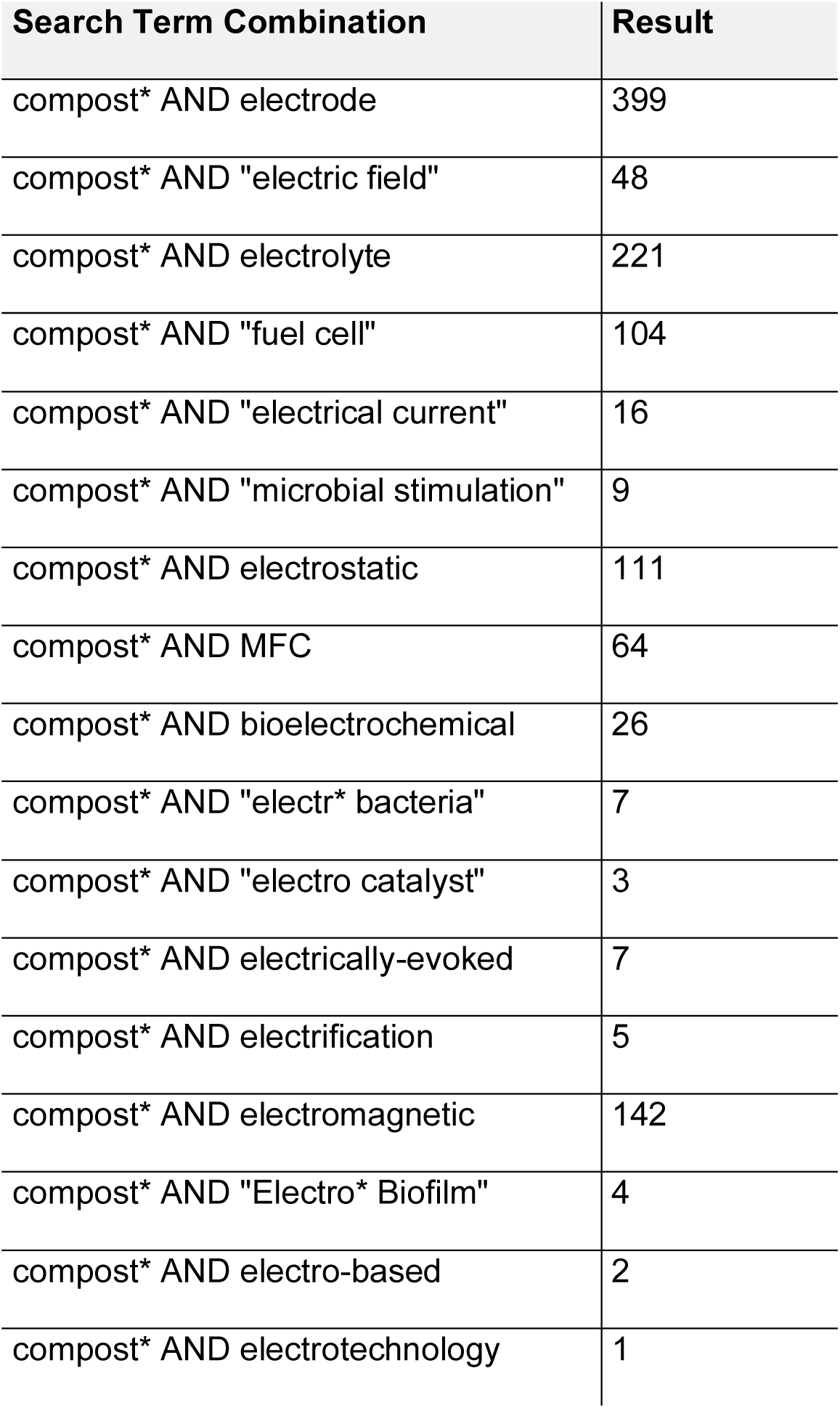
Number of Articles found for each search term before applying exclusion and inclusion criteria.

**Figure 1.**
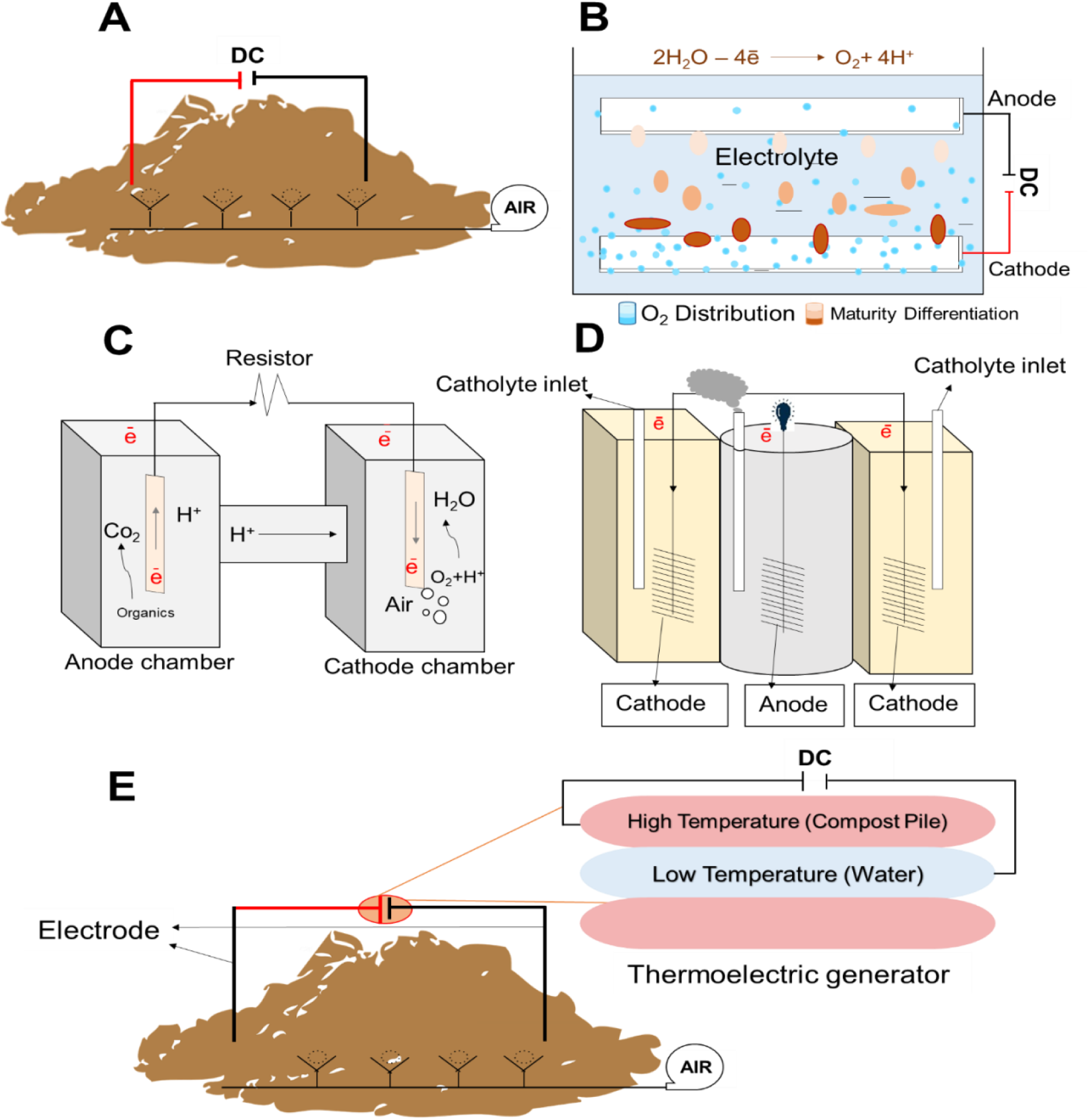
A: Schematic drawing of electric-field assisted aerobic composting (EAAC) system based on Tang et al. (2019); (B) Schematic drawing of electrolytic oxygen aerobic composting (EOAC) system based on Shangguan et al. (2022); (C) Schematic drawing of a microbial fuel cell (MFC) based on Castro et al. (2014); (D) Schematic drawing of the three-chamber bioelectrochemically assisted anaerobic composting (AnC_BE, III_) based on Yu et al. (2018); (E) Thermoelectric generator (TEG) in aerobic composting based on Shangguan et al. (2020).

**Figure 2.**
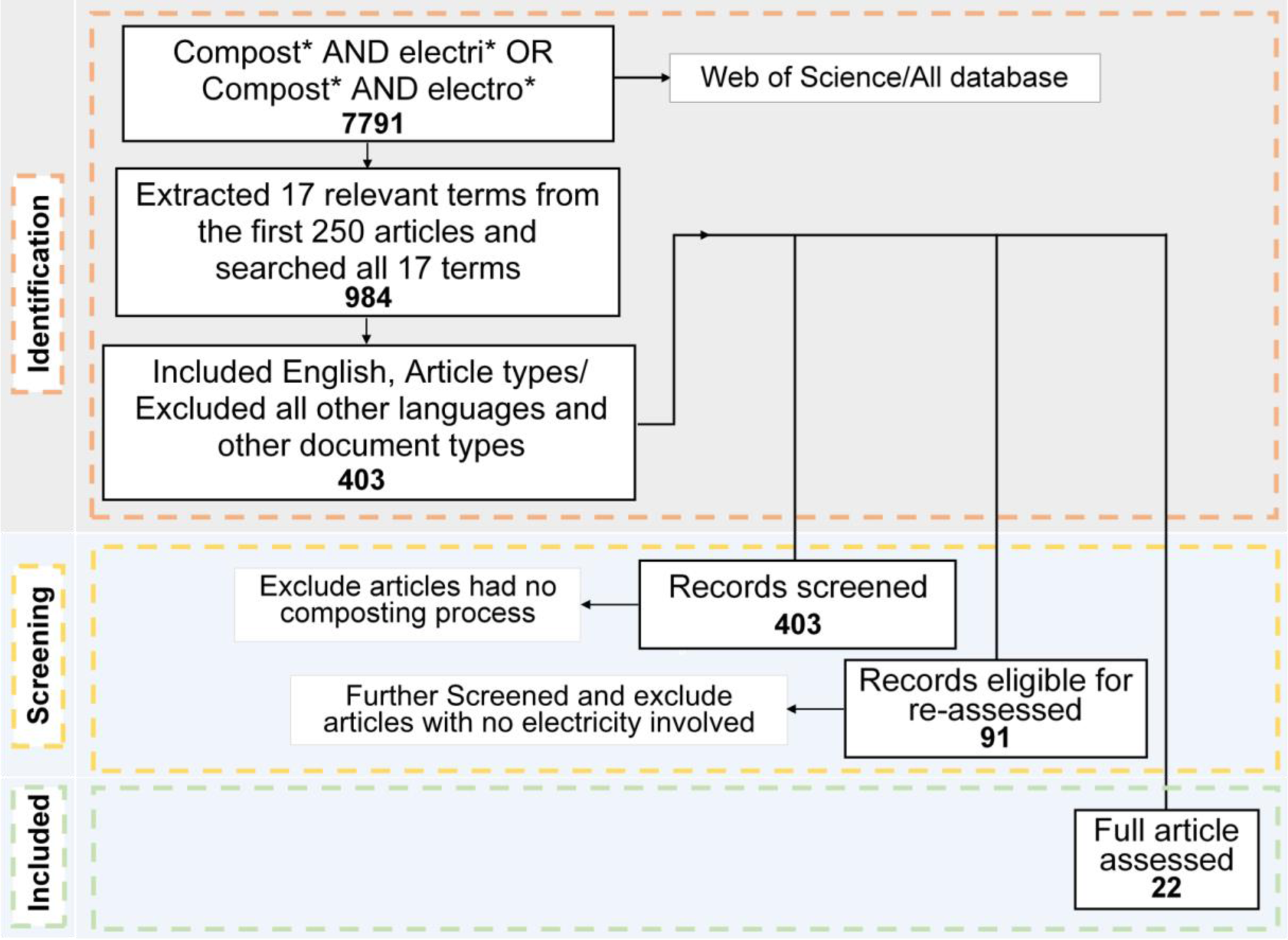
Illustrates the process conducted for the systematic literature review. **Figure illustrations by colors:** 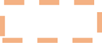: Initial search and broad screening. 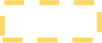: Refined search with specific keywords. 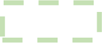: Final screening and selection stage.

**Figure 3.**
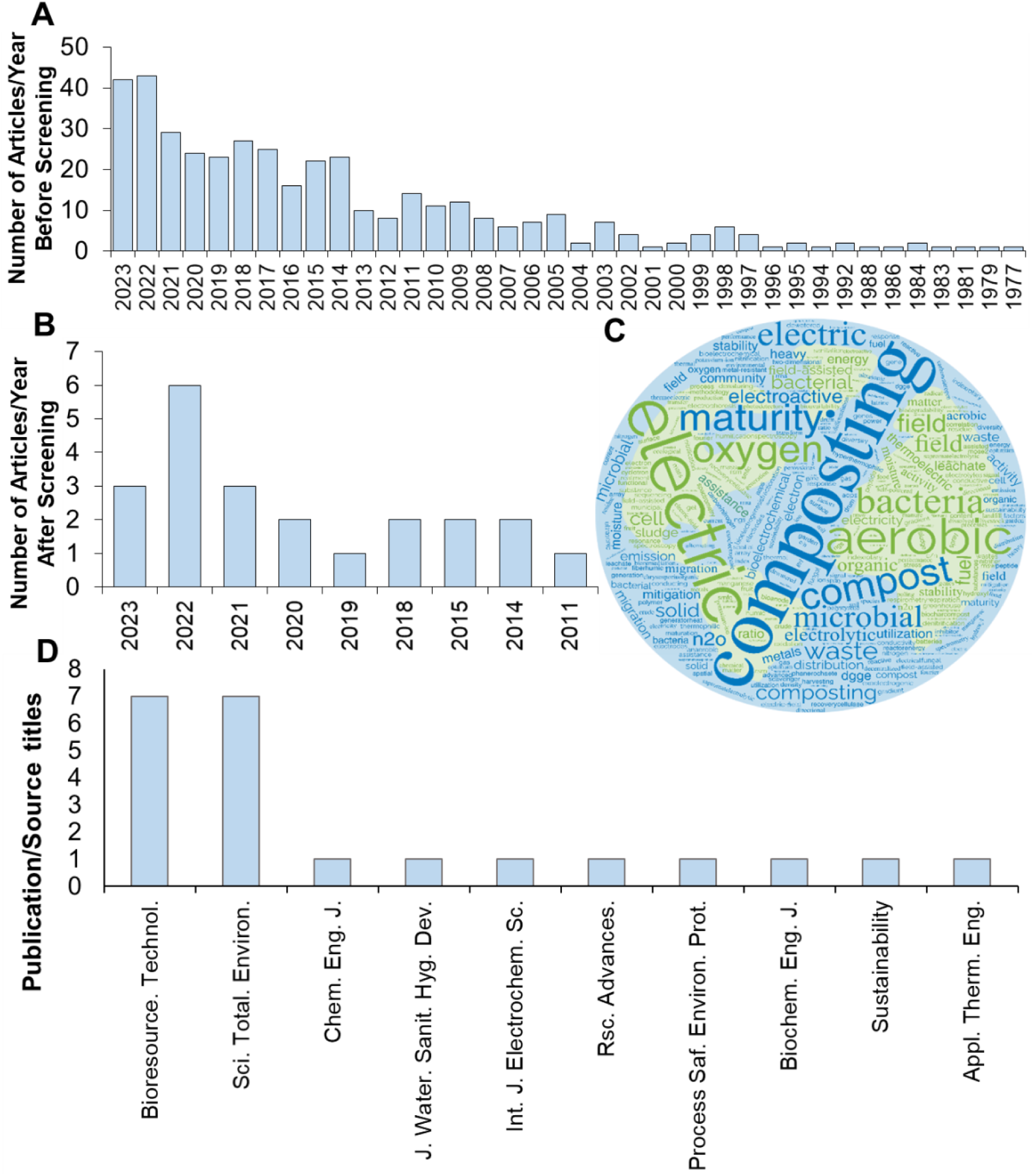
Bibliographic overview. (A) Number of articles/year before screening; (B) Number of articles/year after screening; (C) Word cloud highlighting keywords used in the 24 articles; (D) Publication/source title.

## 3. Results and discussion

In the reviewed articles, four main electro-composting systems including electric-field assisted aerobic composting (EAAC), electrolytic oxygen aerobic composting (EOAC), microbial fuel cells (MFC), and thermoelectric generators (TEG) were covered in more than one article. The remaining system concerns rare cases, for which only one article was found. This includes bioelectrochemically assisted anaerobic composting (AnC_BE, III_).

Seven critical parameters including important materials for the systems setup, operating conditions, temperature evolution, compost maturity, microbial communities, and key features for each system were compared.

### 3.1 Electric-field-assisted Aerobic Composting (EAAC)

The authors noted that several abbreviations have been used in the literature. This includes EAAC for electric-field assisted aerobic, EAC for electric field assisted composting and AEFAC alternating electric field assisted aerobic composting. To make it less confusing, the authors decided to use exclusively the term EAAC. Figure 1A illustrates the concept of electric-field assisted aerobic composting, which incorporates electric fields to enhance composting efficiency and increase microbial activity. EAAC is a new technique that shows promise for boosting compost quality, decreasing emissions of greenhouse gases, and making composting more economically viable.

#### 3.1.1 Materials for the EAAC setup

chicken manure and rice hulls were important in the studies because of their widespread availability and their specific contribution to the composting process as chicken manure provides a high nitrogen content and rice hulls are rich in carbon. To maintain 65-70% moisture, chicken manure, and rice hulls were mixed with dewatered sewage sludge and mature compost in a ratio of 5:2:2:1 (w/w) (Fu et al., 2022). Additionally, in poultry manure, Cao et al. (2021a, 2021b) used sawdust together with poultry manure in a 3.5:1 ratio in a 50 L reactor.

With this ratio, they achieved a C/N ratio of 20, similar to Fu et al., a moisture content of 65%. Another substrate that has been tested in EAAC is food waste, Mi et al. (2023) used food waste mixed with chicken manure, mature compost, and rice husks in a 6:2:1.5:1 (v/v) ratio resulting in 60% moisture content. He et al., (2023) used food waste as well, which they mixed with rice straw and shredded them between 0.5-1 cm in the reactor of 20 L foam boxes. Altogether, the works mentioned above demonstrate the suitability of EAAC for a wide range of substrates. Apart from these conventional substrates, Fu et al. (2021) showed the possibility of further improving EAAC by using supplements. In their specific case, Fu et al. used biochar from rice hulls at 900°C as an additive at a ratio of 9:1 (w/w). Moreover, the other study by Li et al. (2022) utilized chicken, and rice husk at a ratio of 7:2 (w/w) as electric field-assisted composting and conventional composting, and another reactor used chicken, and rice husk including biochar mixed at a ratio of 7:2:0.8 (w/w) as biochar-added electric field-assisted aerobic composting. The biochar was prepared from bamboo at 600°C, crushed, and sieved to 0.5-2 cm.

#### 3.1.2 Design of EAAC composters

although they vary in shape, size, and material, the design of electric-field-assisted aerobic composting (EAAC) systems is similar in most cases. Most authors used a well-insulated reactor containing anode and cathode for electrical stimulation. To give some examples: Tang et al. (2019) used 200 L cylindrical casks with a DC voltage for electric-field-assisted aerobic composting (EAAC), including a stainless-steel sheet (150 cm x 50 cm x 0.3 cm) as positive and graphite (diameter 10 cm, height 60 cm) as negative electrodes. In a similar study, Fu et al. (2022) used 4.5 cm thick foam rectangular reactors with stainless steel plate electrodes. Cao et al, (2021a) used a 50 L bench-scale reactor with graphite as electrodes. Mi et al. (2023) used 150 L cube reactors. A power source is needed to actively stimulate the underlying microbiome. For this, a conventional potentiostat can be used, as in Cao et al. (2022). To allow more diverse configurations, it is also possible to apply a multi-potentiostat as in Li et al. (2022). Although most studies have rather similar approaches, there are some cases, where EAAC is improved due to the application of biochar. For example, Fu et al., (2021) improved the EAAC performance using biochar as a supplement. Usually, direct current is applied. However, in contrast to this, Fu, et al., (2022) have also shown the application of alternating current. Apart from reactor size, shape, and input materials, systems are deviating with respect to the electrode material and as well in the voltage applied. In this regard, He et al. (2023) can be highlighted, who used 2 V DC EAAC reactors with carbon felt and Fe plates as electrodes. In another example, Mi et al. (2023) used array electrodes for Pin-EAC and Flat-EAC configurations, where Pin-EAC referred to the reactors with array electrodes and Flat-EAC referred to the reactors with flat electrodes. Comparing these two types of electrodes, Mi et al. found that using array electrodes greatly raised the composting temperature and enhanced the germination index, resulting in a more successful and faster composting process. In respect to the voltage, there are some variations. Some systems work with low values such as 1 V (Tang et al., 2019) or 2 V (Cao et al., 2021a; Cao et al., 2022, He et al., 2023). Other systems applied higher values of up to 5 V (Tang et al., 2019; Fu et al., 2022) or even 10 V as in Mi et al.,2023.

#### 3.1.3 Operating conditions during EAAC

studies on optimizing composting systems suggest several approaches to increase microbial activity and organic matter breakdown. Tang et al. (2019) took samples from three random places at 20 cm depth to test the physicochemical parameters including moisture content and volatile solids every five days. They experimented with multiple DC voltages (1-5 V) while the maximum temperature was almost the same between 2 V and 5 V and ultimately chose 2V as well as Tang et al. compared non-aerated composting with alternating aeration. A small pump was employed to deliver oxygen. Days 1-13 employed an alternate aeration technique, with the air turned on and off intermittently every 24 hours at a flow rate of 1.5 L/min. During days 14-30, there was no active aeration. On days 10 and 20, the substrates in each reactor were emptied, mixed uniformly, and promptly returned to the reactors. Fu et al. (2021) mixed the biochar with chicken manure, and rice hulls to maintain high moisture levels. Cao et al. (2021a) tested 2 V and 5 V DC voltages, continuous aeration at 0.2 L/min/kg dry weight, and periodic mixing and sampling for 42 days. Mi et al. (2023) suggested online temperature sensors to better keep track of the reactor performance. An interesting approach comes from Cao et al. (2022). To improve the operating conditions, they evaluated different electrolytes. At a voltage of 2 V, they mixed each of the following electrolytes such as FeCl_3_ (5% w/w), KCl (6.9% w/w), and NaHCO_3_ (0.8% w/w) with the composting substrate to mitigate greenhouse gas emissions. As a result, they found that FeCl_3_ was successful in decreasing emissions of ammonia, but it increased the nitrous oxide emissions, illustrating the importance of carefully considering the trade-offs when choosing electrolytes for composting.

#### 3.1.4 Temperature evolution during EAAC

several studies have found an impact of electric fields on the temperature. The alternating electric field-assisted aerobic composting system exhibited more uniform water distribution, resulting in higher initial temperatures (90.85 ± 0.76°C by day 3) compared to conventional aerobic composting (73.95 ± 1.15°C by day 4) and sustained temperatures over 80°C for four days (Fu et al., 2022). Fu et al. (2021) compared EAAC without biochar as well as biochar without electric stimulation. The combination of electric fields and biochar raised the temperature to 71.3°C on day 4, while the controls remained at 68.2°C (EAAC) and 59.8°C (biochar). The combination of EAAC and biochar kept temperatures above 50°C for 16 days. The porosity of biochar enhances air permeability, which helps to elevate the composting temperature. This enabled temperatures up to 77.5°C. Another work that also applied biochar in EAAC achieved 72.6°C and remained above 55°C for more than 7 days, which confirms the positive impact of biochar on EAAC (Li et al., 2022). Another work from Fu et al. was already applied at full scale (Fu et al., 2022). In the resulting high-temperature composting the authors reached 91°C and remained above 80°C for 3 to 5 days (Fu, et al., 2022). Although there are systems that reach high temperatures, there are counterexamples. Low temperatures were observed in studies by Tang et al. and Li et al. In the study from Tang et al., an EAAC system reached just 65°C and it stayed above 55°C for 15 days. However, this was again better than the control. The control showed its temperature peak at 58°C and remained above 55°C for just 10 days (Tang et al., 2019). In the case of Li et al., EAAC piles reached 67.8°C for 10 days, while the control piles just reached 58°C for 5 days (Li et al., 2023). In the study by Tang et al., the EAAC system temperature rose to 68.5°C, while the conventional aerobic composting control peaked at a temperature of 63.1°C, probably due to better organic carbon degradation in the EAAC system (Tang et al., 2020).

#### 3.1.5 Compost maturity in EAAC

compost maturity is an important metric in all composting-related investigations. Compared to conventional composting, EAACs with alternating electric fields increase Humic acid (HA) and fulvic acid (FA) molecules by 429.58 ± 10.95%. These are heterogeneous molecules with a high polymer composition which serve an important role in improving nutrient accumulation and humification in composting. The EAACs have further 65% higher electron accepting and electron donating capabilities, which speeds up lignin oxidation and increases quinones/aromatic compounds. In the same approach, compost maturity was reached in 20 days, with a 28% higher humification index (HI) and a 38% higher germination index (GI) content (Fu et al., 2022). In another study by Tang et al., EAAC together with biochar speeded up the compost maturation by 33%, producing more HA- and FA-like compounds and stabilizing dissolved organic matter (DOM) faster than the respective control (Tang et al., 2019). Mi et al. (2023) used a so-called Pin-EAAC, where Pin-EAAC referred to reactors with array electrodes. They compared the pin electrodes with flat electrodes and found a 13% higher GI for pin electrodes than for flat electrodes. On the other hand, reactors with flat electrodes showed increased humification and a 33% quicker maturity period (Mi et al., 2023). Due to the implementation of EAAC systems, advanced composting technologies, can help to manage waste more sustainably by increasing humification, microbial activity, and composting time.

#### 3.1.6 Microbial community of EAAC

researchers indicate that bacterial communities in EAAC are enriched in *Bacillus*, *Navibacillus*, *Ureibacillus,* and *Thermobifida*. *Ureibacillus* and *Navibacillus* bacteria were the most prevalent in EAACs with alternating electrical fields (52.36% on day 3 and 46.54% on day 41) compared to conventional aerobic composting (2.37% and 12%). High temperatures supported thermophilic bacteria such as *Pseudomonas*, *Ardenticatena*, *Firmicutes*, and *Bacillus* (Fu et al., 2022). In the study by Tang et al. (2019), denitrifying bacteria rose from 3.8% to 8.1% in the control system, but in EAAC, they had decreased to 2.6%. EAAC did not include the main denitrifying microorganisms identified in conventional composting, such as *Nitrospira* and *Denitrobacter*. *Bacillus* and *Alcaligenes,* genera that both contain electroactive species, were observed as the primary denitrifiers in the respective EAAC approach. Another study by Li et al. (2023) described that the abundance of *Pseudomonas and Bacillus* increased to 2.66% and 15.6% respectively in EAAC compared to 1.88% and 4.36% in conventional aerobic composting, while *Acinetobacter’s* abundance gradually decreased after 18 days in both systems.

#### 3.1.7 Major impacts of EAAC

Hyperthermophilic composting (HTC) and EAAC experiments improve efficiency, microbial activity, and environmental impact. Fu et al. (2022) state that compared to conventional HTC, EAAC saves money by aging compost at 80°C using thermophilic microbes as the high temperature speeds up the composting process, shortens the maturation period, and increases microbial activity, which causes faster decomposition and cheaper overall expenses. Tang et al. (2019) discovered that employing EAAC at 2 V direct-current voltage decreased maturation time by 33%, greenhouse gas emissions by 70%, oxygen utilization by 30 ± 9%, and boosted electroactive bacteria abundance by 3.4 times. Fu et al. (2021) found that biochar increases electrical conductivity, lowers methane and nitrous oxide emissions, and accelerates compost maturation by 25%. Moreover, another study reported that EAAC promotes moisture migration and microbial activity. It leads to faster compost maturation, particularly in the cathodic zone (Fu et al., 2022). EAAC inhibits nitrification, increases N_2_O-consuming genes, reduces heavy metal bioavailability by 83.7%, and promotes metal-resistant bacteria (Li et al., 2023). Acidic electrolytes reduce ammonia (NH_3_) emissions by 72.1% while increasing N_2_O emissions due to altered nitrifier activity (Cao et al., 2022). Bioelectrochemical assistance can reduce N_2_O emissions by 28.5% to 75.5%, based on voltage by eliminating ammonia oxidation, highlighting trade-offs (Cao et al., 2021b). Modern composting processes increase efficiency, microbial activity, and environmental impact; however, emission management requires careful consideration.

### 3.2 Electrolytic Oxygen Aerobic Composting (EOAC)

The electrolytic oxygen aerobic composting, shown in Figure 1B, utilizes electrolysis to generate oxygen on-site, ensuring optimal aerobic conditions for composting. Shangguan et al. (2022) and Wei et al. (2022) have both conducted analyses of EOAC, which they compared to conventional composting. They advocate for the use of electrolytic oxygen to enhance the quality and efficacy of composting.

#### 3.2.1 Materials for the setup of EOAC

The study by Wei et al., (2022) used plastic bucket reactors for EOAC. Electrolysis was performed with direct current. Shangguan et al. (2022) employed a raw material ratio of 6:3:1 for chicken manure, mature compost, and rice husk. Electrodes made of a rectangular steel plate and a graphite rod were connected to apply a voltage. Electrodes were positioned at the bottom and the top of the EOAC reactor. Shangguan et al. (2022) focused on oxygen distribution and microbial activity at different depths, whereas Wei et al. (2022) focused on the chemical composition and transformations of DOM.

#### 3.2.2 Design of EOAC composters

both the studies mentioned above-utilized composters with direct electric fields and electrodes to efficiently disperse oxygen. Shangguan et al. (2022) used cylindrical plastic bucket reactors (50 cm diameter, 80 cm tall) with cotton insulation to improve thermal insulation and keep composting temperatures stable. Similarly, Wei et al. (2022) employed rectangular reactors of 50 cm in length, 40 cm in width, and 90 cm in height. Three strategically placed sampling ports at 15 cm, 40 cm, and 65 cm from the bottom enabled for accurate vertical profiling of composting material in the reactors from Wei et al. This design helped in understanding the composting process at various depths and allowed for more accurate monitoring of temperature and other parameters. Although the design is similar to an EAAC, it’s a different process. An EAAC is aiming at an electrical stimulation of microbes. In contrast, an EOAC is aiming at improved oxygenation. However, both concepts overlap in their functionality, as some of the EAAC systems use voltages, which are set high enough to cause electrolysis and thereby oxygen formation. Therefore, the authors of this review suggest to clearer distinction between EAAC and EOAC in the coming research.

#### 3.2.3 Operating conditions during EOAC

Shangguan et al. (2022) applied a voltage of 10 V (DC) to the EOAC reactor. In contrast, the control reactors received adequate aeration but remained without electrical stimulation. Amongst others, the study discovered that oxygen dispersion influences microbial activity and compost maturity at varying depths. Wei et al. (2022) collected compost solid samples from different fractions of the reactor at various depths at various time points for a total duration of 30 days. They investigated chemical processes and transformations. They discovered molecular changes such as microbial activity, chemical transformations, oxygen dispersion, and moisture migration during EOAC, indicating that the bottom layer experienced a higher abundance of common thermophilic bacteria (such as *Cerasibacillus*, *Lactobacillus*, and *Pseudogracilibacillus*), which could encourage compost maturation. This comparison demonstrates the huge impact of operational settings on chemical process parameters and the underlying biology.

#### 3.2.4 Temperature evolution during EOAC

*Shangguan et al. (2022)* found that the composting process was divided into two phases: (1) temperature-raising during the first 6 days; (2) thermophilic phase/humification for the following 24 days. They assessed molecular alterations in the dry organic matter (DOM). They found that low O/C (oxygen-to-carbon) and high H/C (hydrogen-to-carbon) compounds were preferentially decomposed during EOAC. The decomposition in EOAC was higher than in conventional composting controls. During the EOAC process, temperatures rapidly increased to the thermophilic phase (>50 °C) at all three locations measured. The bottom temperature (84.1±1.3 °C) was significantly higher than the middle and top (Wei et al., 2022).

#### 3.2.5 Compost maturity in EOAC

according to Shangguan et al. (2022), higher maturity is linked to effective oxygen distribution and microbial activity, particularly near the pile’s bottom. Wei et al. (2022) found that EOAC increases the breakdown of complex organic molecules and the creation of humus. EOAC promotes compost maturity by physical oxygen dispersion more evenly through the pile of compost and chemical reactions.

#### 3.2.6 Microbial community of EOAC

the study by Shangguan et al. (2022) found that due to the positioning of the electrodes, thermophilic bacteria (*Cerasibacillus*, *Lactobacillus*, and *Pseudogracilibacillus*) were more prevalent at the bottom of the compost. However, this was likely affected also due to better insulation in that area. A second study was published on EOAC, which investigated how the microbial community interacts with DOM and it was discovered that various microbial processes produce chemical changes (Wei et al., 2022). These changes include converting dissolved organic matter into humic substances, decomposition of organic matter, and generating metabolites including alcohols, gasses, and acidic substances.

#### 3.2.7 Major impacts of EOAC

Both, the articles from Shangguan et al. and Wei et al., agree that EOAC composts are faster, that they use higher amounts of oxygen, and that they achieve higher compost quality than conventional composting. According to the first article by Shangguan et al. (2022), optimizing the oxygen distribution can improve compost maturity and as well composting efficiency. The second article from Wei et al. (2022) demonstrated how EOAC alters DOM’s chemical structure due to the decomposition of complex organic compounds like lignin and cellulose into simpler molecules such as sugars and organic acids, it is vital to highlight that, even when reduced lignin improved the compost quality.

### 3.3 Microbial Fuel Cells (MFCs)

Integrating microbial fuel cells (MFCs) with composting is another form of innovative waste management. This section reviews studies on MFC efficiency and the importance of optimizing environmental factors and microbial interactions. These findings suggest that MFCs can alleviate environmental concerns, but it has also been highlighted that they improve sustainability and efficiency. Figure 1C schematically demonstrates the dual functionality of the microbial fuel cell system, which not only treats organic waste but also produces bioelectricity.

#### 3.3.1 Materials for the setup of compost MFCs

Among the different set-ups published, there are similarities, but also differences with respect to material and system characteristics. Wang et al. (2015) used an acrylic chamber and carbon felt as electrodes. Similarly, Nandy et al. (2015) used carbon cloth as electrodes, however, unlike Wang et al., a 10wt% platinum-coated carbon black was utilized as a micro-film electrode. Nandy et al. reported also on the implementation of a proton exchange membrane (PEM) directly in the soil as part of a microbial fuel cell setup. Wang et al. (2015) pretreated their electrode as well but with a different purpose. They soaked the carbon felt in 10% of hydrogen peroxide for 3 hours to stimulate microbial adherence. Wang et al. (2014) utilized cow manure together with 37% of miscellaneous organic matter. Air-dried, pulverized, and blended manure was employed. Eight titanium electrodes with 189 cm² and a 1571 cm² main electrode were employed in the composting tub with a volume of 3.6 m^3^. The compost was prepared from cow excreta, which was mixed with a microbial additive containing *Jiangella* spp., *Actinomadura* spp., *Glycomyces* spp., *Thermobifida fusca*, *Bacillus* spp., *Planifilum* spp., and *Mechercharimyces* spp. in a ratio of 10:1 (Sugimoto et al., 2011). In a quite extraordinary MFC approach, researchers aimed to create a sustainable sanitation solution that could treat human waste and generate electricity Castro et al. (2014). They constructed the MFC by using locally available concrete blocks. This approach not only addressed sanitation challenges but also contributed to waste management and energy generation in the community. As electrodes, they used graphite granules stressing their sustainability and cost-efficiency. Nandy and Wang concentrated on electrochemical efficiency using advanced materials, whereas Castro et al. investigated practical and scalable solutions for resource-constrained places.

#### 3.3.2 Design of MFC composters

several designs were used to integrate MFCs with composting. Nandy et al. (2015) utilized cylindrical single-chamber MFCs with 50 mL anode chambers and aluminum mesh as current collectors. They used hot-pressed membrane electrode assemblies at 120-140 °C and 6.89 MPa pressure to achieve good electrode contact with the proton exchange membrane (PEM) in a compact, efficient design. The electrode membrane assemblies were designed to provide proper contact between the electrodes and the proton exchange membrane (PEM), which is critical for optimal electron transfer and overall efficiency of the microbial fuel cell. According to the authors, such a configuration improves the MFC’s ability to produce electricity by facilitating the flow of protons and electrons throughout the system. To get further insight into the impact of MFCs on chemical parameters, a study by Wang et al. (2015) used solid MFC setups for C/N ratio and moisture content measurements. SMFCs generate electricity using solid organic waste as a substrate, such as agricultural waste, animal manure, and urban waste, instead of wastewater. They applied reactors with different sizes which include smaller chambers with 200 cm³ volume and 18 cm² of the electrode surface, as well as larger chambers with 2000 cm³ of volume and 65 cm² of electrode surface. They were especially interested in analyzing the pH. They discovered that solid MFC’s ideal pH range was between 6 and 8, which greatly enhanced the generation of electricity. In the case of the latrine MFC, Castro et al. (2014) combined composting and urine treatment with an electrical treatment using anoxic anodes and cathodes. Additionally, they included aerobic nitrification chambers. Therefore, the system contained both aerobic and anaerobic chambers. The MFC latrine contained an anode in the anaerobic chamber, which oxidized organic matter and generated energy. The cathode was put in an aerobic chamber, where nitrification occurred, converting ammonium to nitrate for further processing. To create compost, solid human feces were processed separately in the composting toilet and aerobically decomposed by thermophilic bacteria. Both liquid and solid waste were effectively treated by this integrated system, which also produced electricity and compost. The electrodes were composed of graphite granules. In their set-up, Castro et al. composted their solid waste under aerobic conditions, while the liquid waste was treated in the MFC. This integrated technology uses oxygen-exposed composting to treat liquid waste while also producing electricity. In the case of Wang et al. (2014), which was already introduced further above, they constructed a polymethyl methacrylate single-chamber air cathode with a diameter of 120 mm and a height of 120 mm. A carbon mesh anode with a surface of 113 cm² was coated with glass fibers to prevent short circuits. A larger carbon mesh disk with a Pt catalyst cathode generated power. The system was aerated from the top and it maintained anaerobic conditions at the bottom.

#### 3.3.3 Operating conditions in compost MFCs

MFCs in electro-composting work under various conditions to enhance microbial activity and electricity generation. Nandy et al. (2015) boosted microbial activity in vermicompost soil by moistening it with sugarcane bagasse and by adding leaves. After a day of acclimatization, multimeters were used to measure the resulting voltages. Resistors were used to calculate the resulting power and to create polarization curves. The anode chamber received composting chamber effluent, which included anode-respiring bacteria that degraded organic matter and transferred electrons. The nitrification chamber transformed ammonium from urine into nitrate, which the cathode reduced to nitrogen gas (Castro et al., 2014). Wang et al. (2014) investigated moisture, phosphate buffer solution, catalysts, and electrode areas. The researchers tested different PBS concentrations (50, 100, and 200 mM), adjusted moisture content to 60%, 70%, and 80%, compared runs with and without a catalyst, and used 28 cm² and 113 cm² electrode surfaces. They found that the proper moisture, PBS, and catalyst concentration had a significant impact on MFC power output as well as on waste degradation, and the best performance was attained with a moisture content of over 80%, a concentration of 100 mM PBS, and the addition of 0.1 mg Pt cm^-2^ catalyst.

#### 3.3.4 Temperature evolution in compost MFCs

Castro et al. (2014) implemented their system in a tropical climate, which sustained high temperatures of around 32°C. The warm temperature improved microbial activity but resulted also in performance variations due to variations in the external variables including variations in ambient temperature, levels of humidity, and the frequency of latrines utilized by the community. The compost temperature reached 60°C after 13 days. Following this. The temperature dropped to 40°C over the next 12 days. This experiment yielded a steady 0.5 V DC voltage. The changes in electrical current and compost temperature indicated that electron generation and electrical current were related to bacterial activity (Sugimoto et al., 2011). In the work by Wang et al. (2015), they hot-pressed membrane electrode assemblies at 120°C to 140°C to ensure adhesion and component functionality. The authors performed the manufacturing with a slow cool-down phase. This cooling procedure guaranteed that the electrodes remained in good contact with the proton exchange membrane, which was critical for efficient electron transfer and overall performance of the MFC. After cooling, the assemblies were integrated into the MFC setup. These studies show that temperature control, whether in controlled laboratory settings, natural climates, or high-temperature fabrication processes, has the potential to improve the performance of electro-composting.

#### 3.3.5 Maturity in compost MFCs

maturation experiments in compost MFCs were used to demonstrate how maturation processes and conditions influence the overall system performance. Wang et al. (2015) found that solid microbial fuel cells performed best with the following conditions: a C/N ratio of 31.4:1; 60% of moisture content with a power density of 4.6 mW/m². They further described that excessive and low C/N ratios or moisture levels reduced microbial activity, stressing the importance of a well-balanced substrate. Wang et al. (2014) discovered that MFCs operate more effectively when the carbon-to-nitrogen (C/N) ratio is lowered. Among multiple variables, the presence of a catalyst in the cathode showed the biggest effect on the breakdown of organic matter. This catalyst contributes to the MFC’s efficiency by speeding up electron transport. Other parameters, such as moisture content, phosphate buffer solution concentration, and electrode size, had a lower impact. Castro et al. (2014) discovered that the MFC latrine’s performance increased after a year, with power increasing from 0.18 μW to 6.75 μW and resistance lowering to 0.5 kΩ, indicating biofilm formation and microbial community. Sugimoto et al. (2011) found that mature compost had a constant temperature, pH of 8.0, 14.7% water content, 35.7% carbon, 1.9% nitrogen, and a C/N ratio of 19.1. Finally, Nandy et al. (2015) used vermicompost soil matured over one month, and enhanced soil with nutrients and microbial flora in the anode chamber to boost the MFC performance, which provided a mature microbial community and high organic content.

#### 3.3.6 Microbial community of compost MFCs

Nandy et al. (2015) investigated a vermicompost MFC and highlighted the *E. coli* strain CCFM8333, the *Bacillus cereus* strain BUU2, and the *Pseudomonas monteilii* strain CIP104883. According to Nandy et al., these strains enhanced the MFC’s performance by oxidizing organic molecules and transferring electrons. Unfortunately, the other studies on MFCs did not dwell on taxonomy. Wang et al. (2015) discovered that optimal composting’s C/N ratio and moisture increased their solid microbial fuel cells’ microbial activity, but unfortunately, they did not describe related taxa. Castro et al. (2014) discovered that their MFC latrine harbored anaerobic bacteria in the anode chamber which degraded organic matter, while nitrifying bacteria in the nitrification chamber transformed ammonium to nitrate. Denitrification at the cathode converted nitrate to nitrogen gas. The other study by Sugimoto et al. described that the anode received its electrons due to bacterial degradation processes, where changes in voltage, electrical current, and compost temperature were related to the bacterial activity (Sugimoto et al., 2011). Wang et al. (2014) investigated the anode microbial community using DGGE. In their case, increasing the anode area and decreasing PBS concentration increased microbial diversity.

#### 3.3.7 Major impacts of MFC

experimental settings have an impact on the ability of MFCs to create energy and remove waste. Nandy et al. (2015) used MFC with vermicompost soil to eliminate 66% more COD with a maximum power density of 4 mW/m^2^. As a result, the authors were able to power an LED light with their MFC latrine (Castro et al., 2014). According to Wang et al. (2015), optimal composting conditions at a C/N ratio of 31.4:1 and a moisture content of 60% resulted in a maximum power density of 4.6 mW/m². Wang et al. (2014) enhanced the performances of microbial fuel cells by using a platinum (Pt) catalyst in it and produced an output voltage of 544±26 mV and a power density of 349±39 mW/m². The MFC with pt catalyst values were higher compared to non-catalysts. According to Sugimoto et al. (2011), MFCs have a power density of 0.06 mW/m²in composters and 0.07 mW/m²in agricultural fields. The process not only produces electrical energy but also improves soil fertility and lowers carbon emissions through the degradation of organic matter and bacterial activity. These investigations demonstrate that substrate composition, electrode materials, and operating conditions all have an impact on MFC electro-composting efficiency.

### 3.4 Thermoelectric Generators (TEGs)

The thermoelectric generator system, illustrated in Figure 1E, integrates waste treatment with energy recovery, utilizing heat generated from composting operations to produce power. Rodrigues et al. (2018) and Shangguan et al. (2020) explored thermoelectric generators, a novel method for improving waste management’s energy efficiency and sustainability.

#### 3.4.1 Materials for the Setup of TEGs

both the studies mentioned above-utilized aluminum thermoelectric generators, which effectively converted temperature differences into electrical voltage. In a more complex system, Rodrigues et al. (2018) used multiple thermoelectric couplings (n- and p-type materials) encapsulated in ceramics to improve durability and performance. Shangguan et al. (2020) employed a simple design of side-by-side thermoelectric sheets (40 × 40 × 4 mm; TGM-199-1.4-1.5), heat pipes, and cooling fins to cool the TEG’s cold end.

#### 3.4.2 Design of TEG Composters

Rodrigues et al. (2018) employed an electrical insulator to connect multiple thermoelectric couplings in series and thermally parallel. In comparison, the second article shows a stripe-shaped TEG with ten thermoelectric sheets connected in series. Metal sheets conduct heat to the hot end, and a water-cooled aluminum pipe keeps the cold end cool. Additionally, heat-conductive silicone gel allows for system-wide heat conduction (Shangguan et al., 2020).

#### 3.4.3 Operating Conditions during TEG

Rodrigues et al. (2018) employed composting temperature gradients in the cycle of 33 days. This gradient dictates the system’s voltage output, and temperature difference stability influences efficiency regarding electricity generation. The compost (hot end) and water-cooled pipe (cold end) were kept at different temperatures, and in turn, this temperature difference influenced the system’s voltage output, which ranged from 6.9 to 14.4 V. A regulator kept it stable (Shangguan et al., 2020). The studies utilized aerobic composting to achieve high temperatures for thermoelectric conversion.

#### 3.4.4 Temperature Evolution during TEG

although heat was converted into electricity, both the studies on TEG achieved compost temperatures higher than 55°C, which is characteristic for active composting and excellent for TEG operation. Rodrigues et al. (2018) investigated and described in detail temperature profiles in three different stages to document the effects of external temperatures on compost temperature stability throughout time. In contrast, Shangguan et al. (2020) discovered a temperature difference of 20-45 °C between compost and ambient temperature. This temperature gradient increased the voltage output from 6.9 V to 14.4 V.

#### 3.4.5 Compost Maturity in TEG

According to the first article from Rodrigues et al., the TEG implementation helps to stabilize the compost pile temperature, which promotes microbial activity (Rodrigues et al., 2018). The second article used the GI to assess compost maturity. On day 12, the GI was 107%, which increased to 118% by day 15. In contrast to this, the control group had a GI of just 88% on the 15th day (Shangguan et al., 2020).

#### 3.4.6 Microbial Community of TEG

the TEG setup’s regular temperature settings foster a diverse and active microbial community, which is required for effective composting (Rodrigues et al., 2018). Both studies by Rodrigues et al. (2018) and Shangguan et al. (2020) did not perform any taxonomical study on the microbial community.

#### 3.4.7 Major Impacts of TEG

TEG composting results in effective thermal energy recovery, and lower operational costs. Rodrigues et al. (2018) present a cost-effective thermoelectric heat recovery device that utilizes thermal energy generated during composting. Similar to this, Shangguan et al. (2020) emphasize how the TEG system recovers heat from waste. Shangguan et al. named this concept a self-powered EAAC (sp-EAAC). According to Rodrigues et al. and Shangguan et al., the implementation of a TEG increases composting efficiency, reduces costs, and recovers heat from waste, making the process more competitive and sustainable.

### 3.5 Three-chamber bioelectrochemically assisted anaerobic composting (AnC_BE, III_)

The AnC_BE III_ system combines anaerobic composting and electrochemical processes to improve organic waste treatment and bioenergy generation (Yu et al., 2018). The three-chamber bioelectrochemical-assisted anaerobic composting process is depicted in Figure 1D, highlighting its chambered design for effective anaerobic decomposition and energy recovery.

#### 3.5.1 Important Materials for the Setup of AnC_BE, III_

Material improvements make the AnC_BE III_ reactor more functional than its predecessor, the AnC_BE II_. The cathode and anode electrodes in the anodic chamber are titanium wire-twisted graphite fiber brushes (STS4024 K, Toho Tenax, Japan). The anodic and cathodic chambers are separated by two proton exchange membranes (PEMs). The anodic chamber has a water bath and a port at the top for gas collection (Yu et al., 2018).

#### 3.5.2 Design of AnC_BE, III_ Composter

Yu et al. (2018) describe the AnC_BE, III_ composter has a cylindrical anodic chamber (80 mm diameter, 100 mm height, 380 mL volume) and two cubic cathodic chambers (60 mm x 70 mm x 100 mm, 320 mL volume).

#### 3.5.3 Operating Conditions During AnC_BE, III_

The system uses dewatered sludge and anaerobic conditions were maintained by purging the reactors with nitrogen. To sustain a stable cathode potential, the catholyte contained potassium ferricyanide (32.9 g L^-1^) and monopotassium phosphate (27.2 g L^-1^) (Yu et al., 2018).

##### 3.5.1.4 Temperature Evolution During AnC_BE, III_

During AnC_BE, III_, a water bath provided thermal insulation for the anode chamber. This enhances organic matter decomposition and energy generation during composting due to stabilized microbial activity involving bioelectrochemical processes (Yu et al., 2018).

#### 3.5.5 Compost Maturity in AnC_BE, III_

Yu et al. found that the initial TOC degraded dramatically within the first 24 days of testing. The AnC_BE, III_ removed 41.2 ± 0.4% of TOC during the 42-day test, surpassing the AnC_BE, II_ (30.3 ± 0.5%) and AnC results.

#### 3.5.6 Microbial Community of AnC_BE, III_

Yu et al. (2018) disclosed that bioelectrogenesis improved electricity and organic matter degradation. Yu et al. did not provide any taxonomical insights about the microbial population.

#### 3.5.7 Major Impacts of AnC_BE, III_

A novel three-chamber bioelectrochemically assisted anaerobic composting system improves energy generation and organic matter breakdown. It outperforms the two-chambered AnC_BE, II_ in terms of power density (7.0-8.6 W/m³) and TCOD removal (42.3% vs. 32.1%). AnC_BE, III_ technique improves organic solubilization while stabilizing and lowering treated sludge volume.

## 4. Conclusion

The research looks at the four main electro-composting systems, namely EAAC, EOAC, MFC, and TEG, as well as AnC_BE, III_. Through the use of ideal materials, design configurations, and operating conditions, EAAC improves compost quality, decreases emissions of greenhouse gases, and boosts the profitability of the composting process. To keep aerobic conditions and improve composting rate and quality, EOAC uses electrolytic oxygen. Managing environmental factors and microbial interactions is crucial for improving system efficiency in MFCs, which convert organic waste into power. By using thermoelectric phenomena to produce electricity from composting heat, TEGs can recover and sustain energy. The alternative system AnC_BE, III_ is less popular, but it has its uses and benefits. The three-chamber bioelectrochemically assisted anaerobic composting system improves energy generation and makes composting better.

Dominant systems are more efficient, cost-effective, and environmentally friendly than less prominent systems due to their extensive use and better technical integration. While MFCs and TEGs recover energy, EAAC and EOAC are efficient and eco-friendly. The efficiency and scalability of electro-composting systems should be the primary goals of future research. Investigating technological synergies in electro-composting, creating cheaper, more long-lasting materials, and optimizing system designs for different kinds of waste are all part of this. Extensive environmental impact evaluations are necessary for the maintenance of these systems.

By improving composting efficiency, decreasing emissions of greenhouse gases, and enabling energy recovery, electrical technologies promote sustainable waste management. The sustainability and economics of organic waste management could be enhanced through electro - composting, which takes advantage of system strengths while resolving present limitations. To achieve environmental sustainability and economic viability in waste management, electro - composting innovation and research are crucial, as this extensive analysis demonstrates.

**Table 2:**
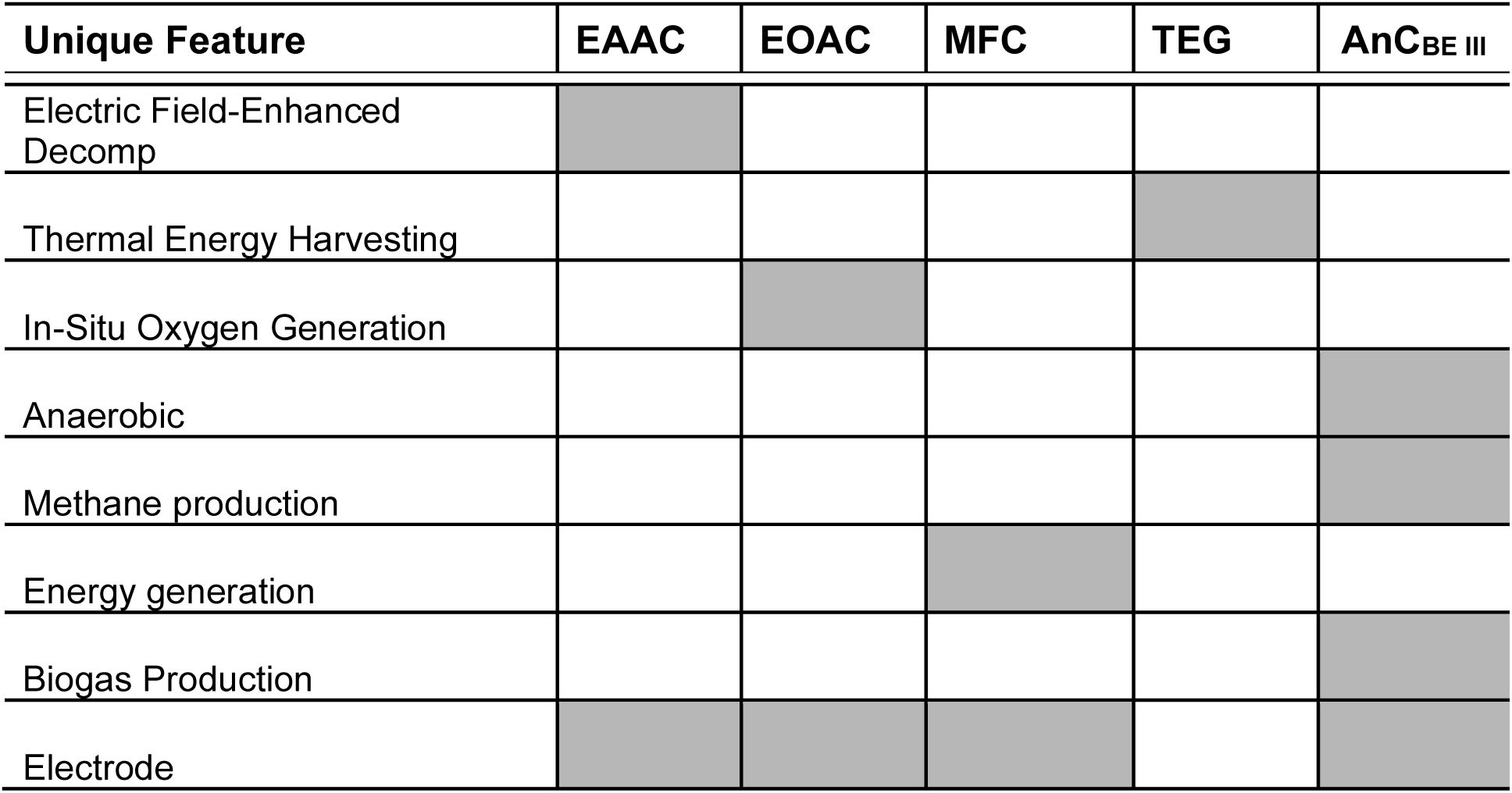
Unique features of electro-composting systems.

## Acknowledgements

The authors are grateful for funding by German Academic Exchange Service (DAAD; PhD grant 57645448 for Ahmad Shabir Hozad).

